# Protein unfolding. Thermodynamic perspectives and unfolding models

**DOI:** 10.1101/2022.11.08.515403

**Authors:** Joachim Seelig, Anna Seelig

## Abstract

Protein unfolding is a dynamic cooperative process with many short-lived intermediates. Cooperativity means that similar molecular elements act dependently on each other. The thermodynamics of protein unfolding can be determined with differential scanning calorimetry (DSC). The measurement of the heat capacity provides the temperature profiles of enthalpy, entropy and free energy. The thermodynamics of protein unfolding is completely determined with these thermodynamic properties. We emphasise the model-independent analysis of the heat capacity. The temperature profiles of enthalpy H(T), entropy S(T) and free energy G(T) can be obtained directly by a numerical integration of C_p_(T). In evaluating different models for protein unfolding. It is essential to simulate all thermodynamic properties, not only the heat capacity. A chemical equilibrium two-state model is a widely used approximation to protein unfolding. The model assumes a chemical equilibrium between only two protein conformations, the native protein (N) and the unfolded protein (U). The model fits the heat capacity C_p_(T) quite well, but fails in simulating the other thermodynamic properties. In this review we propose a modification of the chemical equilibrium two-state model, which removes these inconsistencies. We also propose a new statistical-mechanical two-state model based on a simple, two-parameter partition function Z(T), from which all thermodynamic parameters can be derived. The thermodynamic predictions of the new models are compared to published DSC-experiments obtained with lysozyme, a globular protein, and β-lactoglobulin, a β-barrel protein. Good fits to all thermodynamic properties are obtained. In particular, the models predict a zero free energy for the native protein, which is confirmed experimentally by DSC. This is in contrast to the often-cited chemical equilibrium two-state model, which predict a positive free energy for the native protein. Two-state models use macroscopic fit parameters, the conformational enthalpy and the heat capacity difference between native and unfolded protein. These simulations provide no molecular insight. The review therefore includes a recently published multistate cooperative model based on physicality well-defined molecular parameters only.

## 1. Introduction

Many proteins are only marginally stable at room temperature and can be denatured by heating or cooling. The analysis of protein stability is thus an important problem in developing biological therapeutics. Protein unfolding is a cooperative process with many short-lived intermediates. Cooperativity means that similar molecular elements (e.g. amino acid residues) act dependently on each other. Despite the relevance of cooperativity, protein heat- and cold-denaturation was so far analyzed almost exclusively with a chemical two-state equilibrium [1-9]. This model considers only two types of protein conformations in solution, the native protein (N) and the fully unfolded protein (U). No molecular interactions are specified in a two-state model, which therefore must be classified as non-cooperative. The often-cited chemical equilibrium two-state model as presented, for example, in reference [10] fits the heat capacity quite well. However, it is not consistent with the results obtained with differential scanning calorimetry (DSC) regarding the temperature profiles of enthalpy, entropy and free energy. In this review we therefore present new two-state models, which avoid these difficulties Better agreement with DSC is obtained with an empirical modified chemical equilibrium two-state model, in which all thermodynamic functions are multiplied by the extent of unfolding (Θ_U_(T)-weighted chemical equilibrium two-state model). And even more precise fit of the DSC data s is obtained with a completely different statistical-mechanical two-state model, which is based on rigourous statistical thermodynamics.

The fundamental parameter of all thermodynamic properties of protein unfolding is the heat capacity C_p_(T), which can be measured with differential scanning calorimetry. In this review we first show how the thermodynamic functions enthalpy H(T), entropy S(T) and free energy, G(T) can be obtained by numerical integration of C_p_(T) without the application of any protein unfolding model. We then discuss the standard and the Θ_U_(T)-weighted version of the chemical equilibrium two-state model. Finally, we propose two newstatistical-mechanical models based on rigourous thermodynamic partition functions. The models are compared to DSC measurements of lysozyme and β-lactoglobulin. We emphasise again the model-independent analysis of heat capacity in terms of enthalpy, entropy and free energy. The experimental data are then used to test the different models. The statistical-mechanical models are based on rigorous thermodynamics and provide perfect simulations of all measured thermodynamic properties. The chemical equilibrium two-state model shows discrepancies with respect to entropy and free energy and casts doubt on the physical reality of the postulated positive free energy of the native protein. A modified chemical equilibrium two state model is proposed which corrects most of the insufficiencies. A historical perspective of the chemical equilibrium two-state model can be found in references [7, 8].

## 2. Method – Differential scanning calorimetry (DSC). Model-independent thermodynamic analysis of protein unfolding experiments

Differential scanning calorimetry (DSC) is the method of choice to study the thermodynamic properties of protein unfolding. Details of the DSC method can be found in references [3, 4, 9, 11]. DSC measures the heat capacity C_p_(T) and the relevant literature is focused almost exclusively on the simulation of the heat capacity peak associated with protein unfolding. However, DSC can do more. By numerical integration of the heat capacity the fundamental thermodynamic properties of protein unfolding can be derived. that is [12, 13]

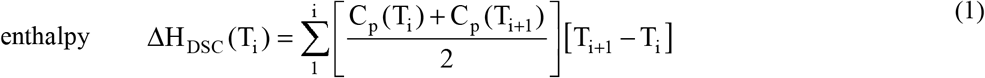

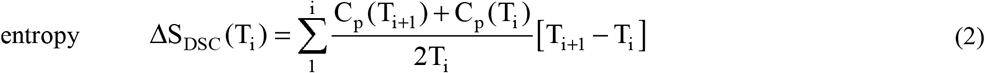

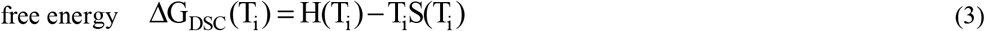

Note that all thermodynamic properties can be evaluated without resorting to a particular unfolding model.

Native proteins have a substantial heat capacity.[14] After unfolding, the heat capacity is even larger. The increase 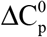, is caused essentially by the binding of additional water molecules [15].

A typical DSC-thermogram is shown in figure 1A. C_p_(T) starts out almost linearly with the basic heat capacity of the native protein. Unfolding gives rise to a sharp heat capacity peak, which is followed by a region of again rather constant heat capacity of the unfolded protein. The choice of the DSC-baseline is important and is handled quite differently in the literature.[9] The subtraction of a sigmoidal baseline is quite common such that native and denatured protein have zero heat capacities [16]. With this correction the further analysis is limited to the conformational change proper. The increase in the basic heat capacity 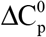 of the unfolded protein is ignored. However, this has been criticized as “it is clear that in considering the energetic characteristics of protein unfolding one has to take into account all energy which is accumulated upon heating and not only the very substantial heat effect associated with gross conformational transitions, that is, all the excess heat effects must be integrated” [11].

**Figure 1.**
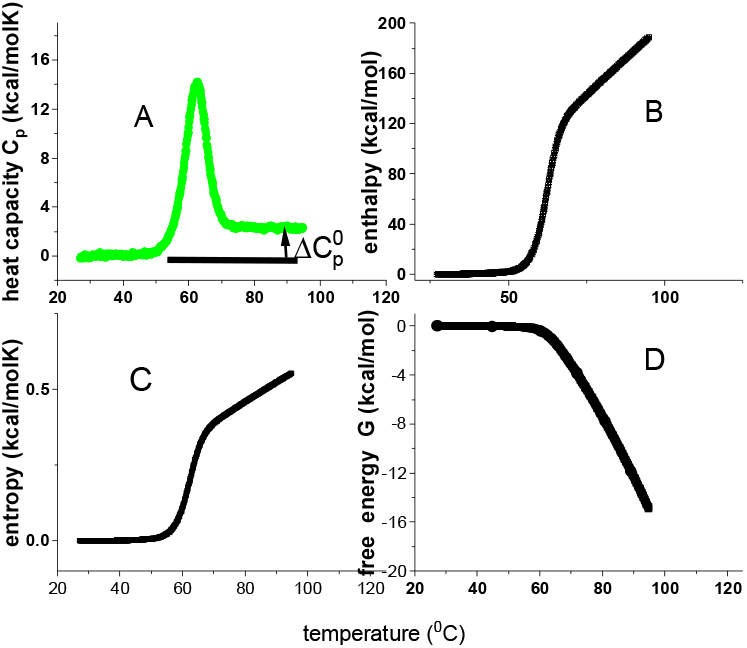
DSC of lysozyme. Model-independent thermodynamic analysis (50 μM, 20% glycine buffer, pH 2.5,). (A) Heat capacity. DSC data (temperature resolution 0.17 °C) taken from reference [17, 18]. (B) Enthalpy ΔH_DSC_(T) (eq.1). (C) Entropy ΔS_DSC_(T) (eq.2). (D) Gibbs free energy ΔG_DSC_(T) (eq.3).

The analysis of DSC experiments as performed in this review is shown in figure 1 for the thermal unfolding of lysozyme [17, 18]. Lysozyme is a 129-residue protein composed of ∼25% α-helix, ∼40% β-structure and ∼35% random coil in solution at room temperature [17]. Upon unfolding, the α-helix is almost completely lost and the random coil content increases to ∼60%. Thermal unfolding occurs in the range of 43 °C < T < 73 °C and is completely reversible. Lysozyme is the classical example to demonstrate two-state unfolding [6, 10, 19].

Figure 1A displays the heat capacity C_p_(T) [17, 18]. The midpoint of unfolding is at T_m_ = 62 °C. Panels 1B -1D show the summation of C_p_(T_i_) according to equations 1 – 3. Due to baseline correction, the basic heat capacity of the native lysozyme is removed and the figure shows the heat capacity of the unfolding transition proper. The heat capacity is hence zero for the native protein according to basic thermodynamics it follows that all thermodynamic properties must also be zero for C_p_ = 0 cal/molK.

As shown in figure 1A, the heat capacity C_p_(T_i_) is a non-linear function of temperature. Consequently, enthalpy H(T) = ∫C_p_ (T)dT, entropy 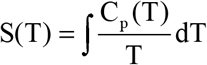 and Gibbs free energy G(T) = H(T) − TS(T) are also non-linear in temperature T. Indeed, enthalpy and entropy display sigmoidal temperature profiles (figs. 1B,1C). The free energy of lysozyme is zero for the native protein, is slightly negative up to the midpoint temperature T_m_, and decreases rapidly beyond T_m_ (fig. 1D).

Figure 1A is typical for the heat capacity profiles of protein unfolding. Surprisingly, the relevant literature reports only heat capacity measurements, but not the corresponding temperature profiles of enthalpy, entropy or free energy, even though these thermodynamic functions become essential in a model-guided analysis.

Unfolding models generally assume baseline-corrected thermograms with a zero heat capacity for the native protein. Equations, 1-3 are not limited to baseline corrected thermograms. Evaluations which include the substantial heat capacity of the native protein found in a recent publication on cooperative protein unfolding [12].

## 3. Theory. Models for protein unfolding

### 3.1. Chemical equilibrium two-state models

Protein unfolding is a cooperative process. Nevertheless, in spite of many short-lived intermediates, protein unfolding is almost exclusively described by a chemical equilibrium between a single native protein conformation (N) and a single denatured molecule (U). The temperature-dependent equilibrium constant is defined as

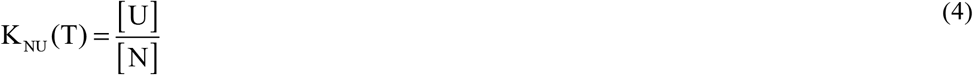

[U] and [N] denote the concentrations of unfolded and native protein, respectively. The temperature dependence of K_NU_(T) is handled differently in different models.

#### 3.1.1. Van’t Hoff enthalpy model

The early version of the two-state model is based on van’t Hoff’s law.[1, 3, 4] The temperature dependence of the chemical equilibrium constant is given by

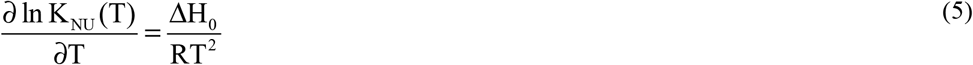

ΔH_0_ is the conformational enthalpy. Integration of equation 5 yields lnK_NU_ (T) = (−ΔH_0_ / RT) + C

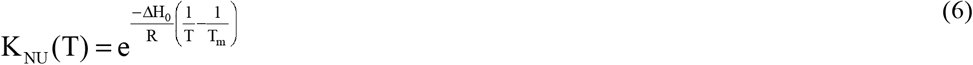

The integration constant C was chosen such that the equilibrium constant is unity at the midpoint temperature of unfolding T_m_, that is, K_NU_(T_m_) = 1. At T_m_, native and unfolded protein occur at equal concentrations.

In the van’t Hoff model the enthalpy is temperature-independent and the unfolded protein has the same basic heat capacity as the native protein.[3, 4, 20]. To account for the experimentally observed increase in the heat capacity of the unfolded protein, the van’t Hoff model was replaced by a more general model with a temperature-dependent enthalpy.

#### 3.1.2. Free energy chemical equilibrium two-state model (“standard model”)

This model is based on the free energy [21, 22]. We follow the common nomenclature [11, 22, 23]. The temperature dependence of the N € U equilibrium is calculated with the free energy ΔG_NU_(T).

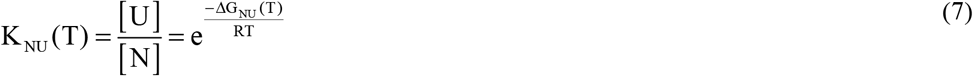

The free energy ΔG_NU_(T) is composed of the enthalpy ΔH_NU_(T) and the entropy ΔS_NU_(T).

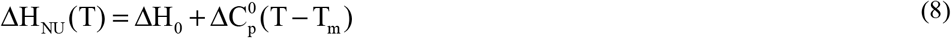

ΔH_0_ is the conformational enthalpy proper and 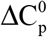 is the increase in heat capacity between the native and the unfolded protein. The entropy S_NU_(T) is defined as

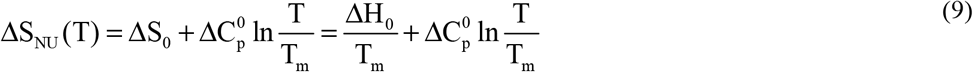

The conformational entropy ΔS_0_ is evaluated by assuming a first-order phase transition (e.g. melting of ice). In such a phase transition the total heat ΔH_0_ is absorbed at a constant. temperature T_m_ and the entropy change is ΔS_0_ = ΔH_0_/T_m_. With the assumption the free energy follows as

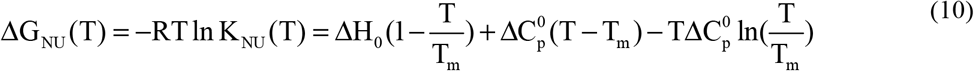

It should be noted however, that in protein unfolding ΔH_0_ is absorbed not at a constant temperature but over a temperature range of 20 - 50 °C.

The extent of unfolding Θ_U_(T) is

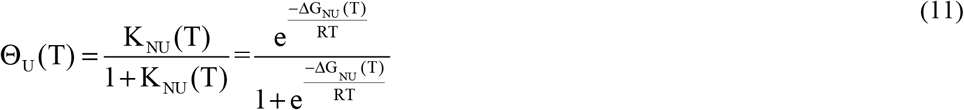

The thermodynamic definition of the heat capacity is C_p_ (T) = (ϑH(T) / ϑT)_p_. The heat capacity of the chemical equilibrium two-state model is

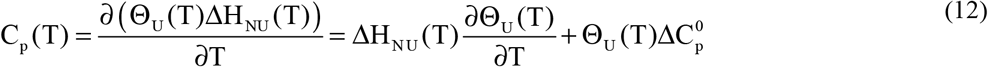

It should be noted that the unfolding enthalpy ΔH_NU_(T) is multiplied with the extent of unfolding Θ_U_(T) to account for the non-linear temperature profile of the heat capacity C_p_(T). Equation 12 is identical to equation 14 in reference [22] and is the hallmark of the standard chemical equilibrium two-state model.

The thermodynamic predictions of this model are shown in figure 2. The enthalpy is a linear function of temperature, the entropy is almost linear, and the free energy ΔG(T) has the approximate shape of an inverted parabola. The native protein has a free energy maximum of 7.51 kcal/mol. At 290 K = 17 °C. However, as the heat capacity is zero at the same temperature this reveals a thermodynamic inconsistency. As shown experimentally in figure 1, a zero heat capacity leads to zero values for all thermodynamic functions. The temperatures for heat and cold unfolding are T_m_ = 63 °C and T_cold_ = -24 °C, respectively. At these temperatures, folded and unfolded protein have the equal concentrations. Cold denaturation may not be feasible experimentally but T_cold_ can be estimated as

**Figure 2.**
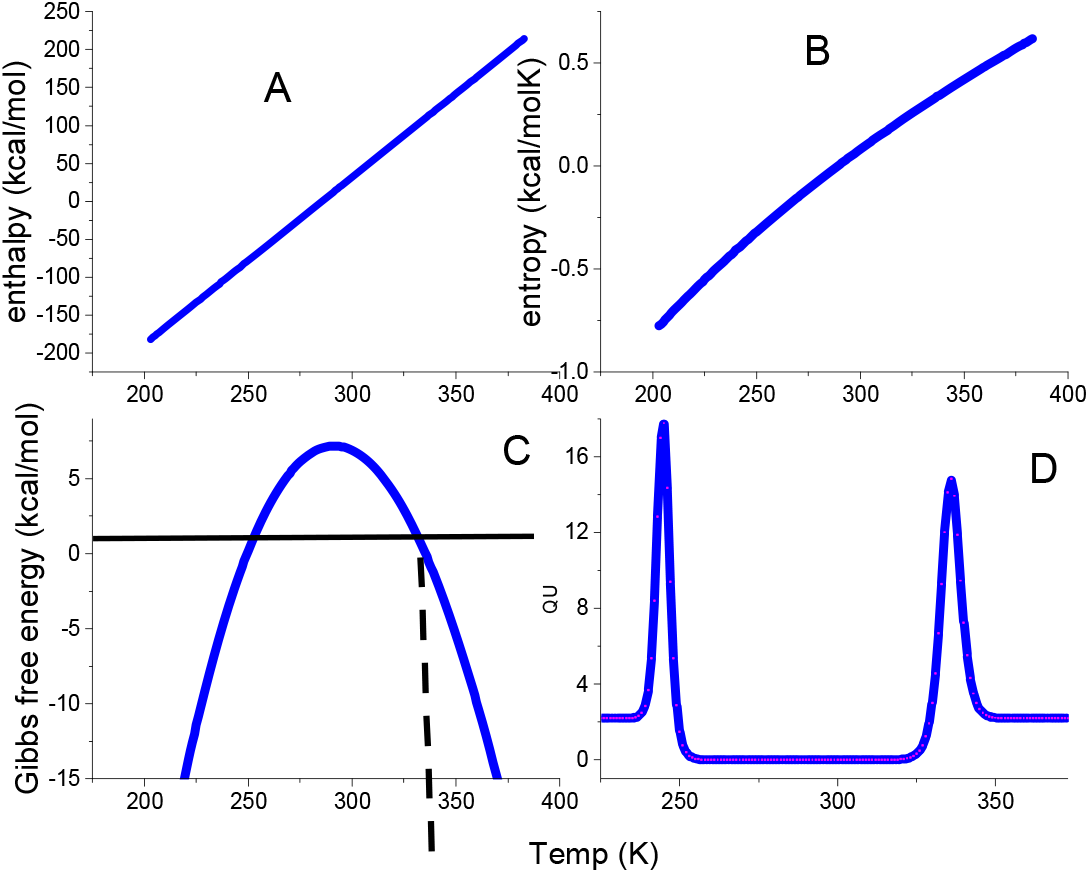
The thermodynamic functions of the standard chemical equilibrium two-state model calculated with ΔH_0_ =110 kcal/mol and 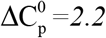 kcal/molK. Dashed vertical line: midpoint temperature T_m_ = 62°C. (A) Enthalpy ΔH_NU_(T). Eq. 8. (B) Entropy ΔS_NU_(T). Eq. 9. (C) Gibbs free energy ΔG_NU_(T). Eq. 10. (D Heat capacity C_p_(T)). Eq. 12.but

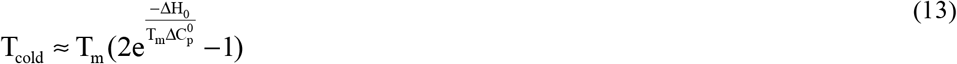

The temperature difference between heat and cold denaturation is

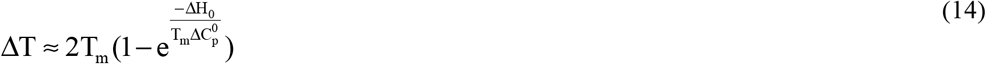

ΔT depends essentially on the ratio 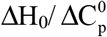. The two parameters have opposite effects. ΔH_0_ increases ΔT, 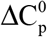 decreases it.

#### 3.1.3. Θ_U_(T)-weighted chemical equilibrium two-state model [24]

The heat capacity of lysozyme in figure 1A shows a non-linear temperature profile. This is taken into account in the chemical equilibrium model by differentiating not ΔH_NU_(T) (which would result in a constant heat capacity, eq. 8), but ΔH_NU_ (T)Θ_U_ (T). As C_p_(T) is non-linear other thermodynamic functions must also be non-linear. The solution is to extend the empirical approach of equation 12 to entropy and free energy. We therefore define a new set of Θ_U_(T)-weighted thermodynamic functions.

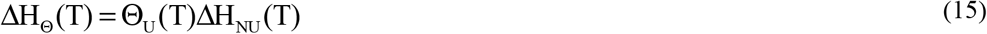

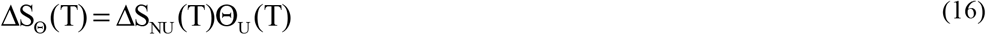

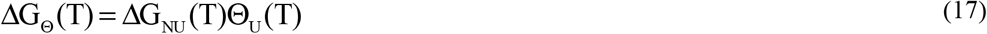

The heat capacity is equation 12 and is not repeated here. The resulting temperature profiles are shown in figure 3.

**Figure 3.**
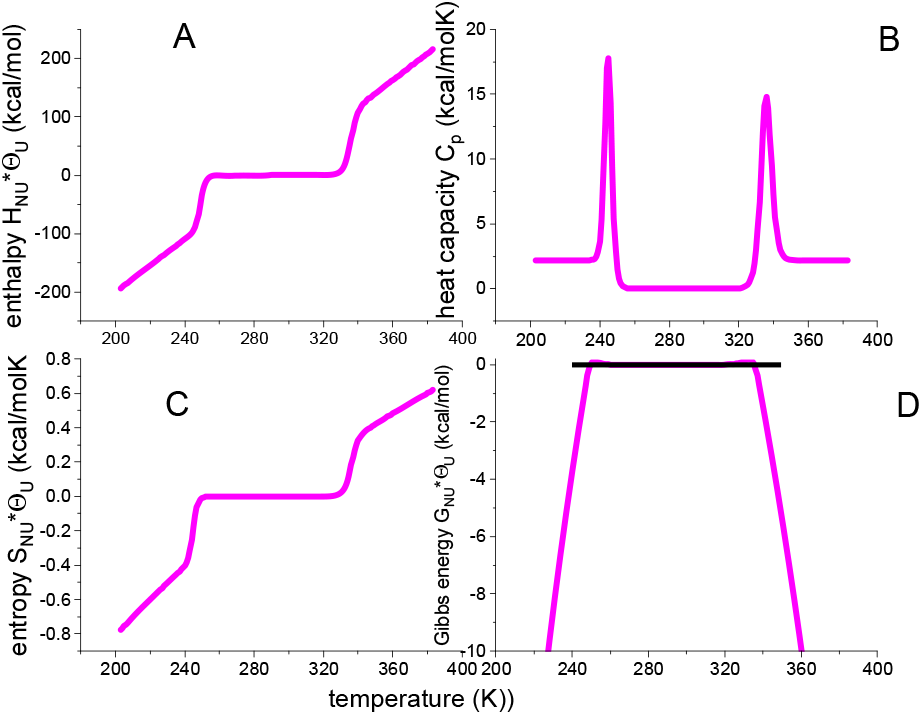
Θ_U_(T)-weighted chemical equilibrium two-state model. (A) Enthalpy (eq. 15). (B). Heat capacity (eq.12) (C) Entropy (eq. 16). (D) Free energy (eq. 17).. Fit parameters: ΔH_0_ =107 kcal/mol, 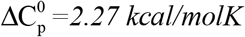

The weighting factor Θ_U_(T) generates sigmoidal temperature profiles for enthalpy and entropy and a trapezoidal profile for the free energy. ΔG_Θ_ (T) of the native protein is zero, which is now consistent with the zero heat capacity. Nevertheless, Θ_U_(T) is an empirical weighting factor, not based on solid thermodynamic reasoning. Indeed, closer inspection of equation 17 reveals residual small positive free energies in the vicinity of the midpoint temperatures T_m_ and T_cold_ (fig. 3D). These are not confirmed by DSC experiment (cf. fig. 7).

### 3.2. Statistical-mechanical models

#### 3.2.1. Partition function Z(T) and thermodynamic properties

Statistical-mechanics provides a rigorous thermodynamic approach to protein unfolding. The heat capacity C_p_(T) is intimately related to the partition function Z(T). Knowledge of the partition function Z(T) then leads to the following thermodynamic relations [25, 26]

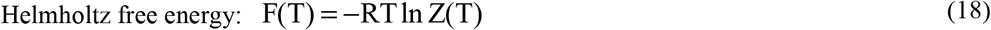

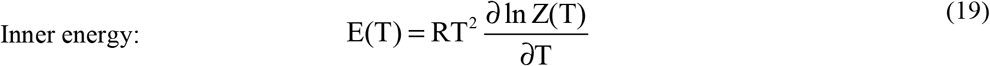

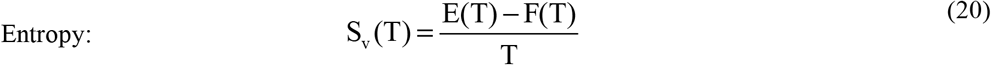

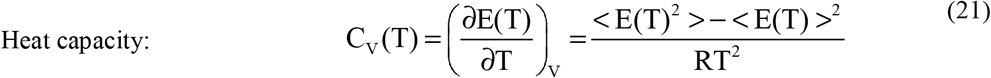

#### 3.2.2. Statistical-mechanical 2-state model [24]

The problem is to find the partition function of a one-component two-state system. Based on the statistics of the linear Ising model as described in reference[27], the following continuous canonical partition function can be defined

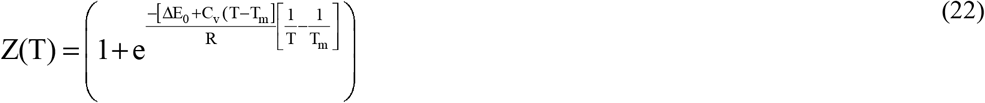

ΔE_0_ is the conformational energy of the unfolded protein. It is temperature-dependent with the heat capacity C_v_. For convenience the partition function is simplified by introducing

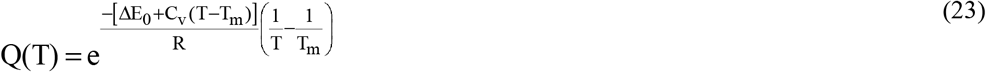

leading to

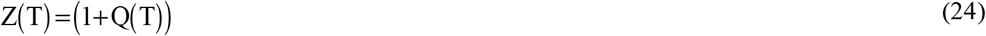

The fraction of unfolded protein Θ_S_(T) is

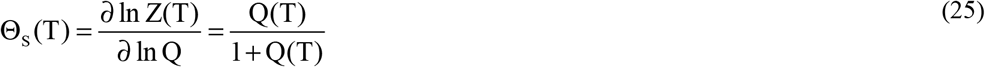

Θ_S_(T) is included for completeness only. It is not needed to calculate thermodynamic functions. The partition function Z(T) suffices to predict all thermodynamic properties. Figure 4 displays the predictions of equations 18-22.

**Figure 4.**
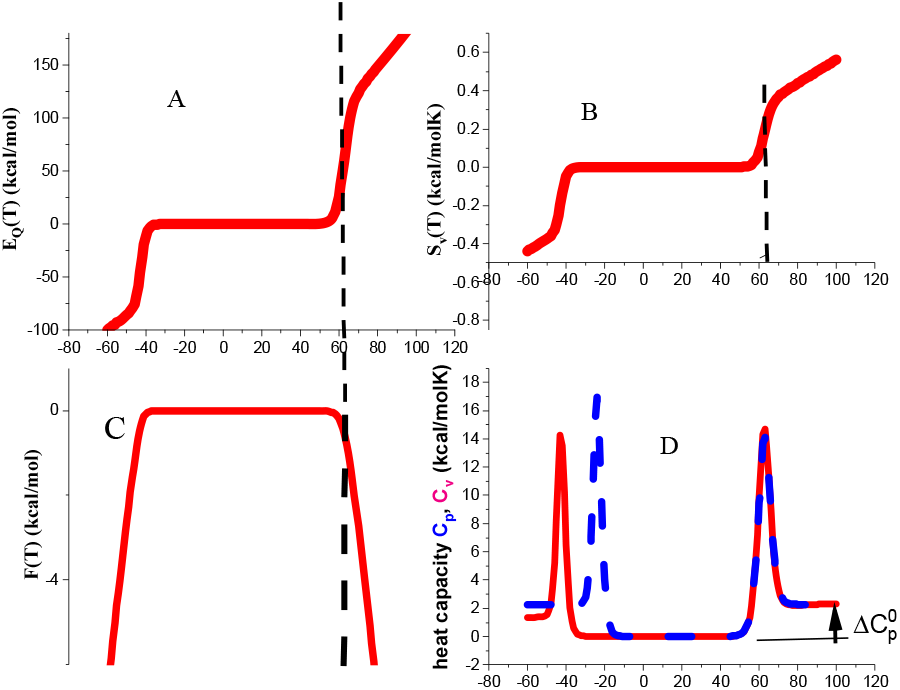
Statistical-mechanical two-state model. Red lines calculated with ΔE_0_ = 110 kcal/mol and C_v_ = 1.05 kcal/molK.. (A) Inner energy ΔE(T). (B) Entropy ΔS_v_(T. (C) Helmholtz energy ΔF(T). (D) Heat capacity C_p_(T). Dashed blue line in panel 4D is the heat capacity calculated with the standard chemical equilibrium two-state model (eq. 12), using, ΔH_0_ =107 kcal/mol, 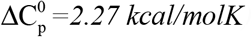 kcal/molK). Dashed vertical lines: midpoint temperature T_m_ = 62°C.

The native protein is the reference state with all thermodynamic functions being zero. In particular, the free energy ΔF(T) is zero at 20°C and becomes slightly negative, but never positive, between T_m_ and T_cold_. ΔF(T) decreases rapidly for temperatures T > T_m_ and T < T_cold_ according to

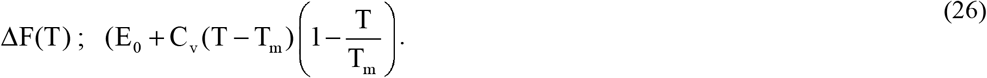

The free energy ΔF(T) of the statistical mechanical two-state model thus also displays a trapezoidal temperature profile. Cold denaturation takes place at

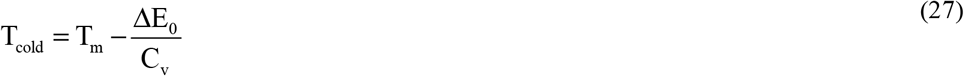

ΔE_0_ and C_v_ have opposite effects on T_cold_. Increasing ΔE_0_ lowers T_cold_, increasing C_v_ leads to an upward shift.

Figure 4D compares the heat capacities predicted by the statistical-mechanical two-state model and the chemical equilibrium two-state model. The high-temperature peaks of the two models overlap precisely but cold denaturation occurs at different temperatures. A discussion of other differences will follow in connection with the protein examples discussed below.

In summary, the statistical mechanical two-state model makes no assumption about the entropy. Also, as mentioned before, no weighting function Θ_U_(T) is needed. to calculate correctly, the thermodynamic properties.

#### 3.2.3. Multistate cooperative unfolding model[12]

The energy of a system with N particles is characterised by its partition function

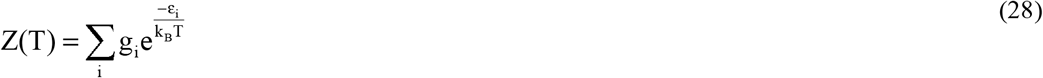

The partition function is the sum of exponential terms (Boltzmann factor) over all energy levels ε_i_, multiplied with their degeneracies g_i_. The partition function determines the thermodynamic properties of the system (eqs. 18 -21) [25, 26]. Here we use the partition function of the multistate cooperative Zimm-Bragg theory, originally developed for the α-helix-to-coil transition of polypeptides [28-30]. Its application to protein unfolding has been discussed recently.[12] The Zimm-Bragg theory has been applied successfully to the unfolding of helical and globular proteins of different structure and size [12, 13, 17, 18, 31-35]. Here we use[12]

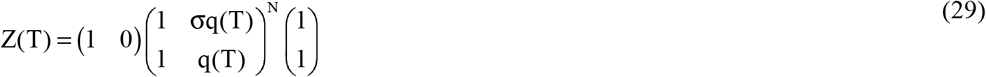

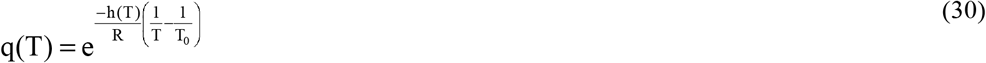

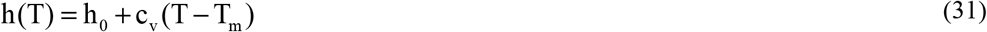

h_0_ is the energy change of the native → unfolded transition of a single amino acid residue. h_0_ is temperature-dependent with the heat capacity c_v_. N is the number of amino acids participating in the transition. The cooperativity parameter σ determines the sharpness of the transition. The σ parameter is typically in the range of 10^−3^-10^−7^. T_0_ is a fit parameter to shift the position of the heat capacity peak. T_0_ is usually close to T_m_. The temperature difference between heat and cold denaturation is 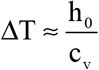.

Figure 5 shows the thermodynamic temperature profiles predicted by equation. 29 in combination with equations 18 - 21. Sigmoidal shapes are predicted for inner energy and entropy, and a trapezoidal shape for the free energy. Figure 5 is very similar to figure 4 of the statistical-mechanical two-state model, but is calculated with molecular parameters only. In particular, the free energy is again zero or negative, but never positive.

**Figure 5.**
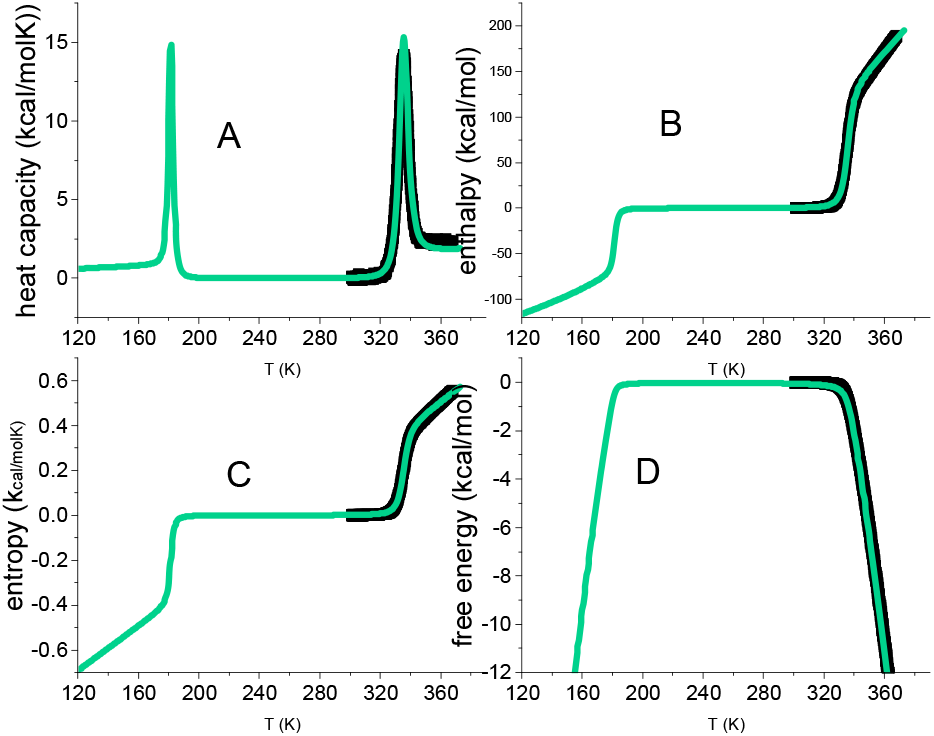
Multistate cooperative model. Green lines calculated with: N = 129 amino acid residues. h_0_ = 900 cal/mol. c_v_ = 7 cal/molK. Cooperativity parameter σ = 5×10^−7^. Black data points: thermal unfolding of lysozyme measured with DSC. Same data as in figure 1.

Figure 5 includes the DSC data of lysozyme heat denaturation (fig. 1). An excellent agreement between theory and experiment is obtained.

The multistate cooperative model describes protein unfolding with molecular parameters of well-defined physical meaning. In contrast, two-state models provide macroscopic parameters.

Equation 29 can be applied to proteins of any size, including large antibodies with ∼∼1200 amino acid residues and unfolding enthalpies of ∼1000 kcal/mol [12].

### 4. Results. DSc experiments compared to protein unfolding models

### 4.1. Lysozyme heat unfolding

Figure 6 compares the experimental date of lysozyme unfolding with two unfolding models. The heat capacity C_p_(T) maximum is at T_m_ = 62 °C and the heat capacity increase is 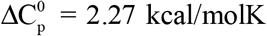, in agreement with literature data of 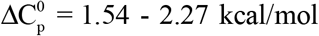 [6, 10, 15, 17, 19, 36, 37]. The red lines in figure. 6 represent the statistical-mechanical two-state model and were calculated with ΔE_0_ = 110 kcal/mol and C_v_ = 1.05 kcal/molK. A perfect fit of the experimental data is achieved.

**Figure 6.**
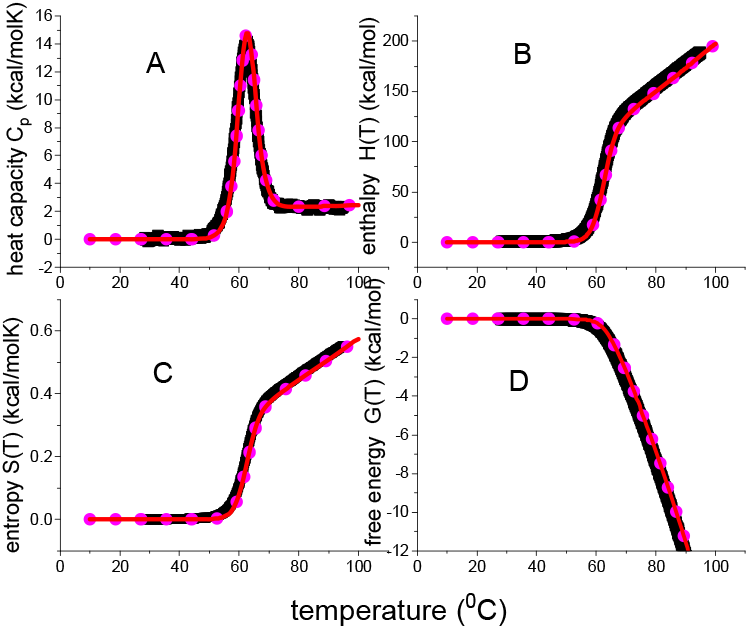
Analysis of lysozyme heat unfolding with 2-state models. Black data points: DSC data of figure 1. Red lines: statistical-mechanical two-state model (ΔE_0_ = 110 kcal/mol and C_v_ = 1.05 kcal/molK). Magenta dotted line: weighted chemical two-state model (ΔH_0_ =107 kcal/mol, 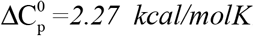).(A) Heat capacity. (B) Enthalpy. (C) Entropy. (D) Free energy.

The magenta dotted lines represent the weighted chemical equilibrium two-state model. The agreement with the experimental data is also very good. However, a small difference to the experimental data is observed near the midpoint of unfolding. Figure 7 displays an an enlarged view of this region. DSC reports a zero free energy for the native lysozyme. The free energy becomes immediately negative upon unfolding. This result is correctly reproduced by the statistical-mechanical two-state model. In contrast, the Θ_U_(T)-weighted chemical equilibrium two-state model (eq. 17) predicts spikes of positive free energy at temperatures just before the midpoints of unfolding. While these spikes are small and of no practical importance, they signify a thermodynamic discrepancy. Of course, the difference between experiment and model would be much larger if the parabolic free energy profile (eq. 10, fig 2C) would be included.

**Figure 7.**
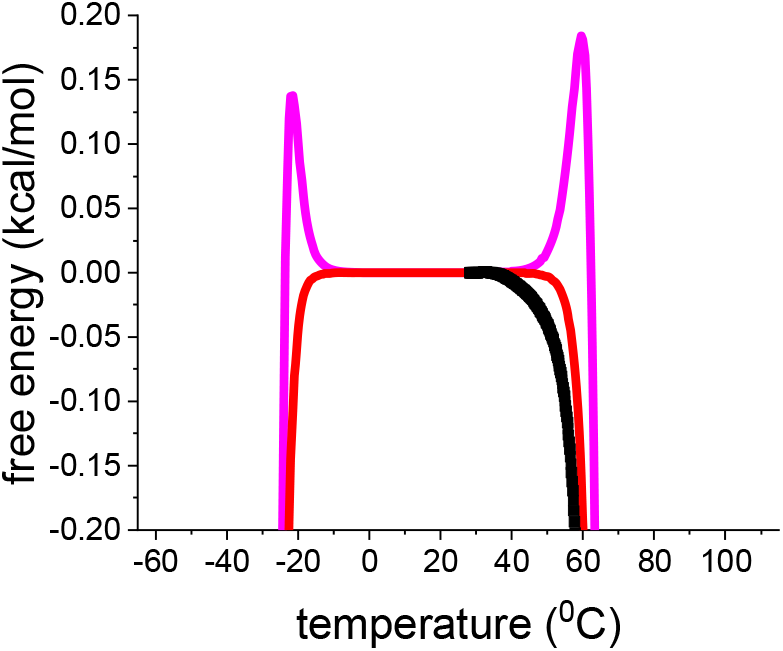
Enlarged view of the free energy. Black data points: DSCresults for lysozyme heat unfolding. Red line: statistical mechanical two-state model (ΔE_0_ = 110 kcal/mol, C_v_ = 1.05 kcal/mol). Magenta line: Θ_U_(T)-weighted chemical equilibrium two-state model (eq. 17, H_0_ = 1107 kcal/mol, 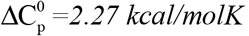).

The multi-state cooperative model as described in figure 5 also yields a perfect fit of the lysozyme DSC experiment [12].

In summary, three different models provide a good to excellent description of lysozyme DSC unfolding. At the midpoint temperature T_m_ all three models predict the extent of unfolding exactly as Θ_U_ = 1/2. Native and unfolded protein occur at equal concentrations. Nevertheless, the free energy is not zero but DSC reports ΔG_DSC_(T_m_)= -0.756 kcal/mol. Indeed, a negative free energy is intuitively plausible as the protein is partially denatured at T_m_. This result is supported by two theoretical models. The statistical-mechanical two-state model yields ΔF(T_m_) = -0.462 kcal/mol, the multistate cooperative model yields ΔF(T_m_)= -0.855 kcal/mol. In contrast, the Θ_U_(T)-weighted chemical equilibrium two-state model predicts ΔG_Θ_(T_m_) = 0 kcal/mol.

### 4.2. β-lactoglobulin. Cold and heat denaturation

Bovine β-lactoglobulin (MW 18.4 kDa, 162 aa) folds up into an 8-stranded, antiparallel β-barrel with a 3-turn α-helix on the outer surface. β-Lactoglobulin in buffer without urea displays only heat denaturation (black squares in fig. 8A, data taken from fig.1 of reference [38]). Unfolding takes place between 55 °C < T < 96 °C with the C_p_(T) maximum at 78 °C. The unfolding enthalpy is ΔH_DSC_ = 74.5 kcal/mol (cf. [38], table 1, 0 M urea), the entropy ΔS_DSC_ = 0.213 kcal/molK, and ΔH_DSC_/ΔS_DSC_ = 349K = 76°C, consistent with the C_p_(T) maximum.

**Figure 8.**
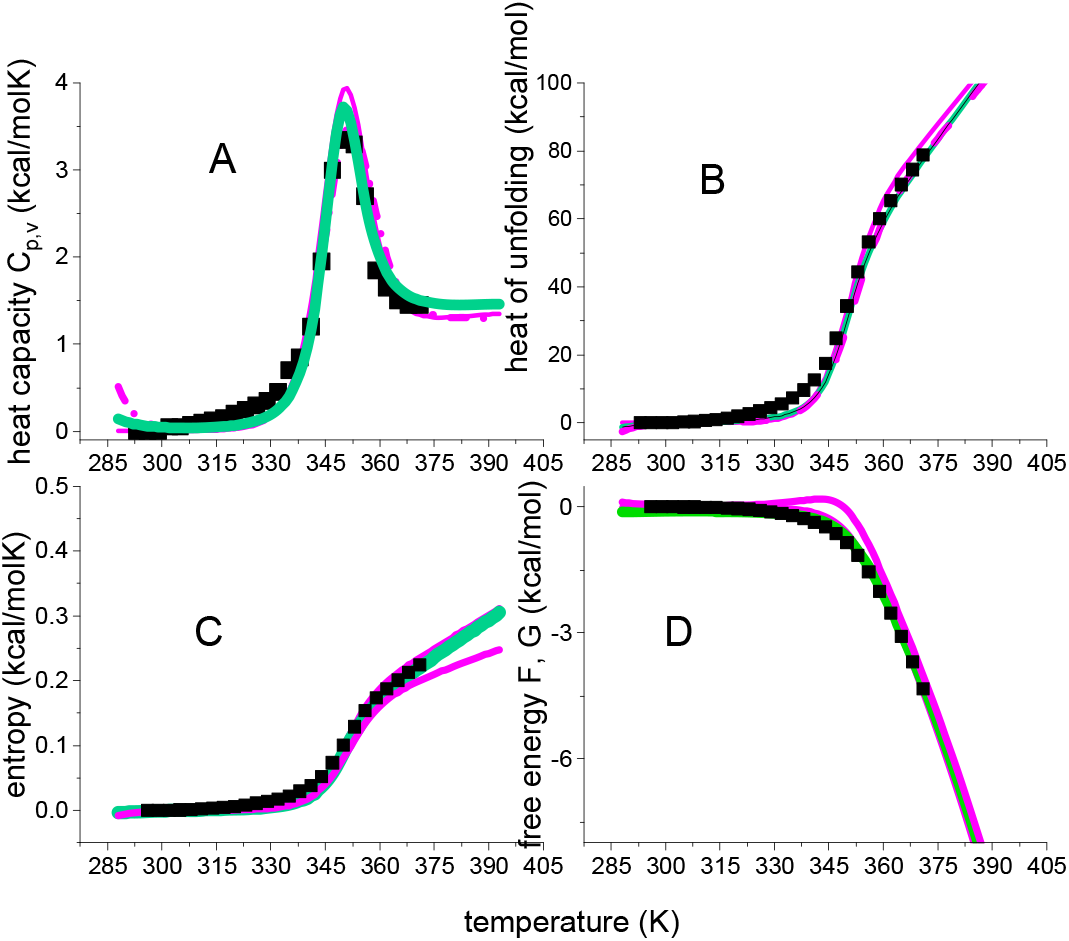
DSC of β-lactoglobulin in 0.1 M KCl/HCl, pH 2.0 buffer. Black data point in in panel A are taken from reference[38](fig. 1). Magenta lines: Θ_U_(T)-weighted chemical equilibrium two-state model (ΔH_0_ = 50 kcal/mol, 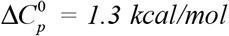). Red lines: statistical-mechanical two-state model (ΔE_0_ = 55 kcal/mol, C_v_ = 0.6 kcal/molK) Green lines: multistate cooperative model (h_0_ = 380 cal/mol, c_v_ = 3 cal/molK, σ = 7×10^65^, N = 160). (A) Heat capacity. (B) Unfolding enthalpy. (C) Unfolding entropy). (D) Free energy of unfolding.

The Θ_U_(T)-weighted chemical equilibrium model, the statistical-mechanical two-state model, and the the multistate cooperative model overlap completely. as far as the heat capacity is concerned. On the other hand, some small differences are observed for enthalpy, entropy and free energy. The most conspicuous difference is seen for the free energy. The Θ_U_(T)-weighted chemical equilibrium model predicts a small peak of positive free energy for the native protein, which is not confirmed by the DSC experiment.

The conformational enthalpy is ΔH_0_ ≌ ΔE_0_ = 50 -55 kcal/mol, which is small for a protein with 162 amino acid residues. Likewise, the molecular enthalpy parameter h_0_ = 380 cal/mol is also small compared to the typical 900-1300 cal/mol of globular proteins.[17] The molecular origin of the small unfolding enthalpy is of β-lactoglobulin can be traced back to its extensive β-structure content. The enthalpy h_0β_ for the reaction β-structure → disordered amino acid was measured as h_0β_ = 230 cal/mol in a membrane environment [39].

A protein can also be unfolded by cooling. Cold denaturation usually occurs at subzero temperatures but can be shifted to above 0 °C by high concentrations of chemical denaturant or extreme pH values. All models discussed above, predict cold denaturation, provided the heat capacities 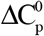 or C_v_ are non-zero. In fact, the temperature difference between heat and cold denaturation depends strictly on the ratio of conformational enthalpy/heat capacity.

Only a few DSC experiments showing at least partial cold denaturation are available. One of the best example is DSC-unfolding of β-lactoglobulin in 2.0 M urea solution (figure 9).

**Figure 9.**
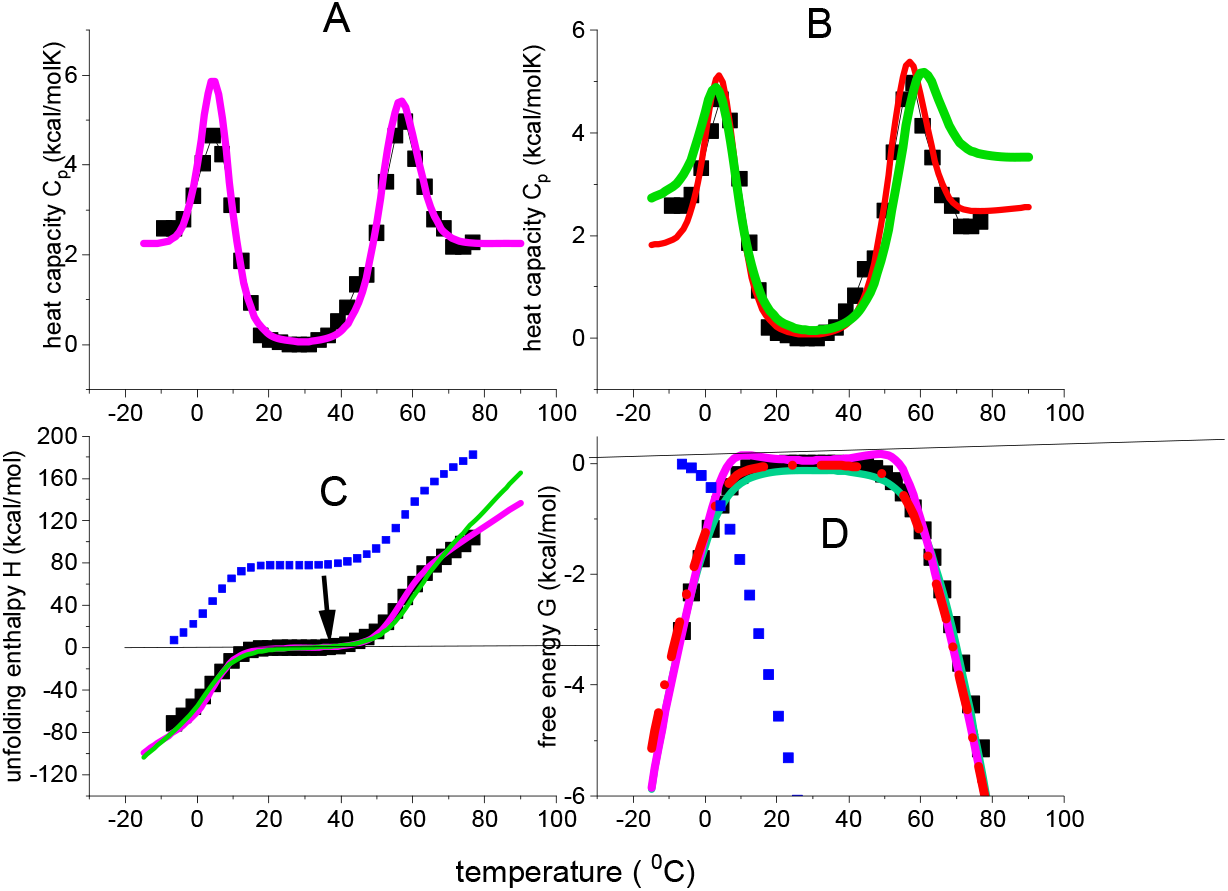
Thermal folding and unfolding of β-lactoglobulin in 2.0 M urea solution. Magenta lines: Θ_U_(T)-weighted chemical equilibrium model. ΔH_0_ = 56 kcal/mol;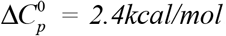 Red lines: statistical-mechanical two-state model. ΔE_0_ = 55 kcal/mol; C_v_ = 1.2 kcal/molK; Green line: multistate cooperative model. h_0_=0.58 kcal/mol, c_v_ = 13 cal/molK, σ = 6×10^−5^, N = 80. (A) DSC heat capacity data taken from reference [38]. Simulation with the Θ_U_(T)-weighted chemical equilibrium model. (B) Same DSC data as in panel A. Simulations with the statisticall models (C) Enthalpy ΔH(T)_DSC_. Integration of the C_p_(T) data according to equation 1 generates the blue data points. The data are then shifted by -78.3 kcal/mol, the enthalpy released upon cold denaturation, resulting in the black data points. This scale shift assigns a zero enthalpy to the native protein. (D) Free energy. Evaluation of C_p_(T) according to equations 1-3 leads to the blue data points. A related scale shifts for the entropy as for the enthalpy (not shown) and recalculation of the free energy results in the black data points. The free energy of the native protein is now zero. Cold and heat denaturation generate negative free energies (for more more details see reference [24]).

The DSC data are taken from figure 2 of reference [38]. The simulation of C_p_(T) is shown in figure 9A for the Θ_U_(T)-weighted chemical equilibrium model and in figure 9B for the two statistical models. The DSC experiment begins at -9°C where the protein is in a disordered state. Heating induces a disord →order transition with a heat capacity maximum at 4°C, a folding enthalpy ΔH_DSC_ = 78.3 kcal/mol and an entropy change of ΔS_DSC_ = 0.282 kcal/molK. The ratio ΔH_DSC_/ΔS_DSC_ is 277K = 4°C, consistent with the heat capacity maximum.

At ∼18-30 °C the protein is in the native-like conformation. Cooling reverses the before mentioned process and returns a disordered conformation with a simultaneous release -78 kcal/mol (cf.fig. 9C).The heat capacity peak at 4°C is hence the mirror image of cold denaturation (see fig. 2 in [38]. Heating β-lactoglobulin above 30°C destroys the native structure. The order → disorder transition has a heat capacity maximum at 57°C, ΔH_DSC_ = 104 kcal/mol and ΔS_DSC_ = 0.312 kcal/molK. The ratio ΔH_DSC_/ΔS_DSC_ of heat denaturation is 333K = 60 °C.

β-Lactoglobulin is less stable in urea solution as the midpoint temperature T_m_ is shifted from 78 °C to 57 °C. Such a decrease in temperature is common in chemical denaturants. However, it is usually associated with a decrease in enthalpy, not an increase [34, 35, 40]. In the present case the enthalpy increases by ∼40%, the entropy by ∼50%, resulting, in turn, in a 20°C downshift of T_m_.

The DSC data were analysed with the Θ_U_(T)-weighted chemical equilibrium model and the two statistical models. The conformational enthalpy ΔH_0_ is equal to the inner energy ΔE_0_ with ΔH_0_ ≈ ΔE_0_ = 56 kcal/mol. These parameters are also identical to those obtained for β-lactoglobulin in buffer. In contrast, the heat capacities 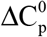, C_v_ and c_v_ are 2 – 3 times large, most likely due to the binding of urea molecules.

The three models discussed above, provide good simulation of all experimental data. In particular, they reproduce the trapezoidal temperature profile of the free energy (black squares in fig. 9D). The native protein has a zero heat capacity and, in turn, a zero free energy. Unfolding leads to negative free energies, both for heat and cold denaturation. The two statistical model exactly the produce this profile. The Θ_U_(T)-weighted chemical equilibrium two-state model displays small positive peaks in the vicinity of T_m_ and T_cold_, which are not supported by DSC

## 5. Conclusions

Under equilibrium conditions protein stability is determined by the midpoint of heat denaturation, by the temperature difference between heat and cold denaturation, and by the width and cooperativity of the unfolding transitions. These parameters are intimately connected to the thermodynamic properties of the system. The thermodynamics of protein unfolding is completely characterised by the temperature profiles of enthalpy, entropy and free energy. The building stone of these thermodynamic properties is the heat capacity C_p_(T), which can be measured precisely with differential scanning calorimetry. In this review we have emphasised the almost completely ignored concept of the direct and model-independent evaluation of the heat capacity in terms of the thermodynamic functions H(T), S(T) and G(T). The evaluation is straightforward and simple. It is hence quite surprising why this approach is not considered in the relevant literature.

Thermodynamic unfolding models should predict not only the heat capacity C_p_(T), but also the complete set of thermodynamic functions. The DSC experiment reveals a sigmoidal temperature profiles for enthalpy and entropy and a trapezoidal profile for the free energy. Focusing on the unfolding transition proper, the heat capacity of the native protein is zero and all thermodynamic functions are equally zero. No positive free energy is measured for the native protein, which is in contrast to the prediction of the often-cited chemical equilibrium two-state model. Experimental temperature profiles were shown for the heat-induced unfolding of the globular protein lysozyme and for the heat and cold denaturation of the β-barrel protein β-lactoglobulin. The experimental results are compared to the predictions of four different models, that is, two chemical equilibrium two-state models and two statistical mechanical models. All four models described the heat capacity equally well. The popular chemical equilibrium two state model fails however in the simulation of the thermodynamic temperature profiles. This model was therefore modified into the Θ_U_(T)-weighted chemical equilibrium model by multiplying the thermodynamic functions with the extent of unfolding. This is is an empirical approach which fits all thermodynamic data quite well, but displays a small discrepancy to the experimental results in the vicinity of the unfolding transition. The two statistical-mechanical models have a rigourous thermodynamic foundation and avoid this difficulty. They provide the best simulations of the experimental data.

Two-state models are non-cooperative approximations to a cooperative multistate protein folding € unfolding equilibrium. They describe protein unfolding in terms of two macroscopic parameters, the conformational enthalpy ΔH_0_ ; ΔE_0_ and the heat capacity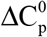: 2C_v_.. In contrast, the multistate cooperative unfolding model uses molecular parameters, that is, the enthalpy h_0_ per amino acid residue, the heat capacity c_v_, the cooperativity parameter σ, and N, the number of participating amino acid residues. The multistate cooperative model can be applied to proteins of any length, e.g. antibodies with 1200 amino acid residues and unfolding enthalpies of 1000 kcal/mol.[12, 35]. In contrast, two-state unfolding models are best suited for unfolding enthalpy is of 50-200 kcal/mol, typically found for small proteins only.

Protein unfolding is characterised by large enthalpies and entropies but a small free energy (enthalpy-entropy compensation). As discussed in detail by comparing lysozyme and lactoglobulin. The free energy is not a good criterion to judge protein stability. Better parameters are defined above. The-beaded chemical equilibrium two state model, the statistical-mechanical two state model and the multistate cooperative model provide quantitative thermodynamic interpretations of these parameters.

